# Elevated plasma cholesterol causally improves sepsis outcome by promoting hepatic metabolic reprogramming

**DOI:** 10.1101/2025.10.22.683940

**Authors:** Qian Wang, Jianyao Xue, Ling Guo, Dan Hao, Misa Ito, Rianna Reese, Bin Huang, Congqing Wu, Xiang-An Li

## Abstract

**Rationale:** Sepsis is a life-threatening condition with high mortality and limited therapeutic options. While hypocholesterolemia is associated with poor outcomes, the causal role of cholesterol and its underlying mechanisms remain unclear.

**Objectives:** To determine whether elevated plasma cholesterol improves sepsis survival and to elucidate the mechanistic basis of this effect.

**Methods:** We analyzed septic patients from the MIMIC-IV database. Total cholesterol levels were compared between 28-day survivors and non-survivors using Wilcoxon rank-sum tests. Cox proportional hazards regression assessed the association between cholesterol levels and mortality, adjusting for age, sex, race, SOFA score, and Charlson comorbidity index. To establish causality, C57BL/6J mice were randomized to receive either a high-cholesterol diet (HCD) or regular diet (RD) for three days prior to cecal ligation and puncture (CLP) to induce sepsis.

**Measurements and Main Results:** Survivors had significantly higher cholesterol levels than non-survivors (median, 135 vs. 126 mg/dL, p < 0.001). High cholesterol (≥ 133 mg/dL) was independently associated with reduced 28-day mortality (adjusted HR = 0.80, 95% CI: 0.67–0.95, p = 0.012). In mice, HCD elevated plasma cholesterol and significantly improved survival (52.5% to 90%), independent of immune modulation.Hepatic transcriptomics revealed metabolic reprogramming, including upregulation of oxidative phosphorylation and glutathione-mediated antioxidant pathways, and suppression of endoplasmic reticulum proteostasis. Notably, inhibition of mitochondrial respiration abolished the survival benefit.

**Conclusions:** Elevated plasma cholesterol causally improves sepsis outcomes by promoting hepatic metabolic reprogramming. These findings offer mechanistic insight into clinical observations and suggest hepatic bioenergetics as a novel therapeutic target in sepsis.

## Introduction

Sepsis is a life-threatening condition arising from a dysregulated host response to infection, leading to organ dysfunction and high mortality rates (1, 2). Despite extensive efforts to target the inflammatory cascade—once considered the primary driver of sepsis—therapeutic interventions have yielded only modest survival benefits (3). This underscores the urgent need to deepen our mechanistic understanding of sepsis and to identify novel therapeutic strategies.

Cholesterol, a vital component of cellular membranes, plays critical roles in maintaining membrane integrity, modulating immune responses, and regulating cellular metabolism (4). Hypocholesterolemia is frequently observed in septic patients and correlates with increased mortality (5-7). Interestingly, the “obesity paradox”—whereby obese individuals exhibit lower sepsis-related mortality—suggests that elevated lipid levels, including cholesterol, may exert protective effects (8-11). While clinical observations implicate plasma cholesterol as a key determinant of sepsis outcomes, the mechanistic basis for this association remains poorly understood. Moreover, prior studies have been limited by small sample sizes (ranging from 17 to 502) and insufficient adjustment for confounding variables (5).

To address these gaps, we integrated clinical data analysis from the MIMIC-IV database with mechanistic investigations in a murine model of sepsis. Our findings reveal that lower plasma cholesterol levels are significantly associated with increased mortality, and that mice fed a high-cholesterol diet (HCD) for three days are markedly protected from cecal ligation and puncture (CLP)-induced sepsis. Mechanistic studies further demonstrate that HCD feeding promotes metabolic reprogramming characterized by enhanced oxidative phosphorylation, glutathione-mediated antioxidant defenses, and suppression of endoplasmic reticulum proteostasis. These insights suggest that cholesterol may act as a critical modulator of host resilience during severe infection, offering new avenues for therapeutic intervention through metabolic pathway targeting in sepsis.

### Experimental procedures

#### Materials and Methods

Materials are listed in the Supplemental Major Resources Table.

#### Retrospective study

The retrospective study utilized the Medical Information Mart for Intensive Care IV (MIMIC-IV) version 3.1, a publicly accessible database containing de-identified health-related data from patients admitted to intensive care units at the Beth Israel Deaconess Medical Center between 2008 and 2022. We loaded the MIMIC-IV dataset in the original CSV file format into DuckDB v1.3.2. We adopted many SQL code from MIMIC GitHub repo (https://github.com/MIT-LCP/mimic-code) to generate derived tables for data analysis in R v4.4.3. The first total cholesterol measurement within the first 24 hours of ICU admission was used for analysis. If ICU data were unavailable, the first total cholesterol measurement obtained during the same hospital stay was used instead. A flowchart outlining the data extraction and patient selection process is provided in Supplemental Fig. 1. All code related to data analysis in the study will be publicly available upon publication.

#### Mice and diet

C57BL/6J (B6) mice were purchased from Jackson Laboratory. Both male and female mice were used in experiments. B6 mice were randomized to feed on a high cholesterol diet (HCD) (TD. 88051, 7.5% cocoa butter, 15.8% fat, 1.25% cholesterol, 0.5% sodium cholate) or a regular-control diet (RD) (2918 Teklad Irradiated Global 18% Protein Rodent Diet, 4.5% fat, 0.022% cholesterol). Diets were administered for 3 days prior to procedures and maintained post-operatively.

#### Rotenone Treatment

Rotenone (Sigma-Aldrich, R8875) was dissolved in ethanol (90 mg/100 ml). This solution was evenly sprayed onto 200g of HCD and allowed to dry thoroughly for one week prior to feeding.

#### CLP-induced sepsis model

CLP was performed on about 3-month-old mice as previously described (12). The cecum was ligated by a 4.0 silk ligature at a half distance of the cecum and punctured twice with a 23G needle. Mice were monitored for survival over a 7-day period.

#### Biochemical assays

Blood was collected from the abdominal aorta at 0, 4, 20, and 44 hours post-CLP. Serum was separated for biochemical assays. The cholesterol kit was from Wako. Serum cytokines were analyzed by Eve Technologies using Mouse Cytokine Array /Chemokine Array 31-Plex (MD31). Corticosterone was measured with an ENZO Life Science kit. Blood glucose levels were measured with a glucose measurement kit (Glucose Meter Kit, Contour) or glucose measurement strips (Glucose test strips, Contour).

#### Coomassie Blue staining of serum Albumin

Serum samples collected 4 hours after CLP were used for albumin detection. Each sample was diluted 1:40 in distilled water, and 10 µL of the diluted serum was loaded per well onto a 10% SDS-PAGE. Following electrophoresis, gels were stained with Coomassie Brilliant Blue to visualize albumin bands. After staining, gels were destained until a clear background was achieved. Albumin band intensity was assessed either visually or quantified using densitometric analysis with ImageJ software.

#### Bacterial load

Blood, peritoneal fluid, spleen, and liver from CLP mice were homogenized and diluted (20, 200, and 2000 times with dH2O), and 100 μL of the dilution was plated on an LB-Agar plate and incubated at 37°C for 24h, and then the number of clones was counted.

#### Flow cytometry analysis of leukocyte recruitment and splenic immunocytes

Leukocyte recruitment to the peritoneal cavity and immunocyte profiling in the spleen were assessed by flow cytometry. Mice were sacrificed at 4 h, 20 h, and 44 h post-CLP. To collect peritoneal cells, 5 mL of PBS was injected into the peritoneal cavity, and the peritoneal fluid was harvested. Total cell numbers were counted, and 1×10^6^ cells were used for staining. Splenic immunocytes were analyzed from HCD and RD mice before CLP and at 20 h post-CLP. After red blood cell lysis, 1×10^6^ splenocytes were prepared for flow cytometry. In both protocols, cells were blocked with CD16/32 antibody to eliminate nonspecific Fc receptor binding. Peritoneal cells were stained with Ly-6C-FITC, Ly-6G-PE, CD11b-PerCP-5.5, and CD45-APC. Splenocytes were stained with B220-FITC, CD3-PE/Cy7, and CD11b-PerCP/Cy5.5. All staining was performed for 30 minutes at 4°C, followed by washing with FACS buffer. Samples were analyzed using a BD LSRII Flow Cytometer.

#### Statistical Analysis

Survival data were analyzed using the Log-Rank test and Kaplan-Meier plots. Comparisons between two groups were performed using a two-tailed Student’s t-test, while comparisons among multiple groups were analyzed using two-way ANOVA. Statistical analyses were conducted using GraphPad software. Data are presented as mean ± SEM, and differences were considered statistically significant at p < 0.05.

## Results

### Low plasma cholesterol concentration correlates with a poor prognosis in septic patients

To better understand the relationship between plasma cholesterol levels and sepsis outcomes, we analyzed data from 2787 sepsis patients (sepsis-3 criteria (1)) with total cholesterol measurements in the MIMIC-IV v3.1 database (13). Baseline characteristics of survivors and non-survivors are summarized in Supplemental Table 1. Survivors had significantly higher median total cholesterol levels compared with non-survivors (135 mg/dL vs 126 mg/dL, p < 0.001), demonstrating a significant inverse association between total cholesterol levels and 28-day mortality (**Fig. 1A**). Patients with higher cholesterol levels showed improved survival outcomes (log-rank, p = 0.00035), with an adjusted (age, gender, race, SOFA score within the first 24 hours, and Charlson comorbidity index) hazard ratio of 0.80 (95% CI: 0.67-0.95, p = 0.012) for high versus low cholesterol groups (**Fig. 1B**). This finding supports the notion that low plasma cholesterol concentrations are a risk factor for sepsis death.

**Fig. 1.**
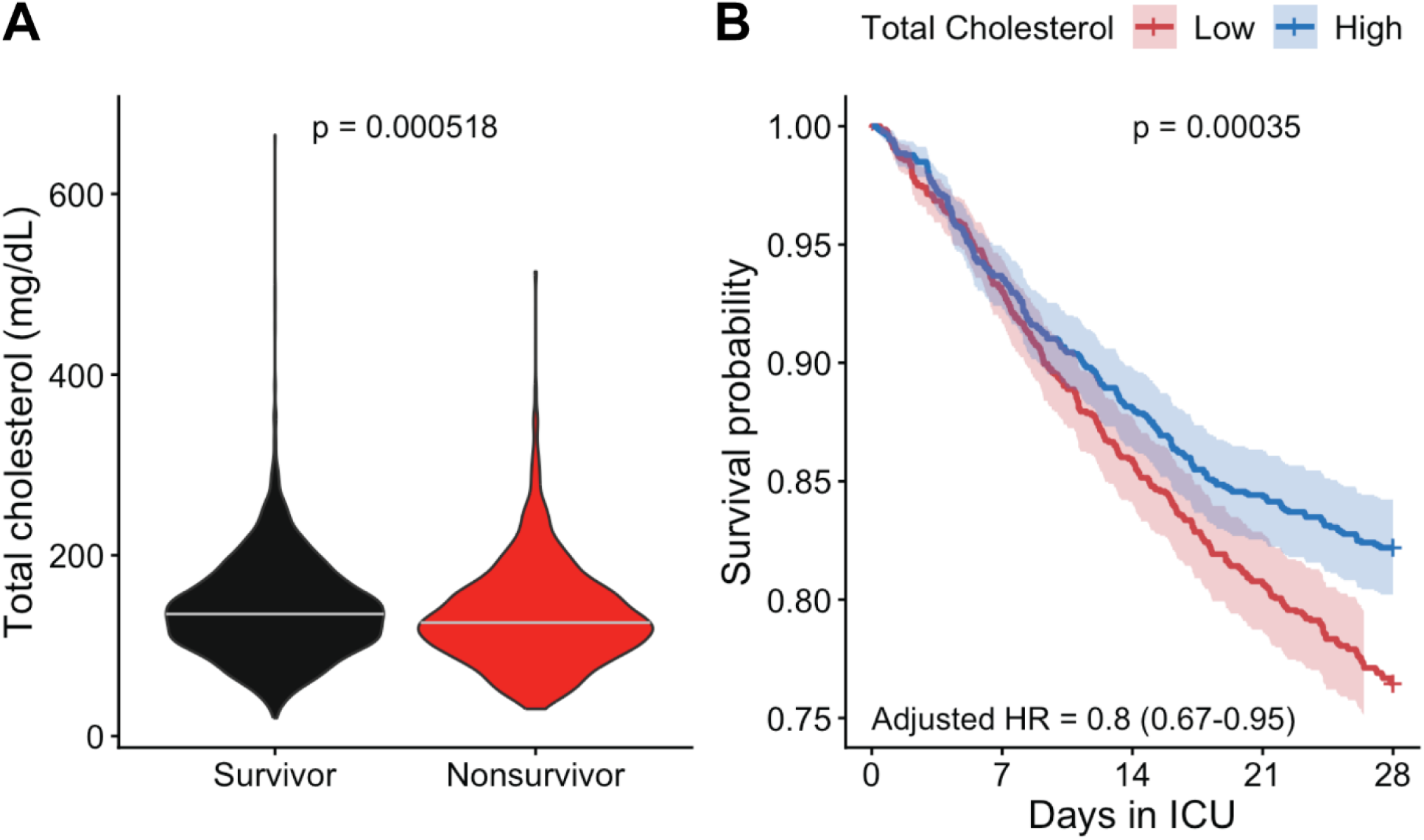
Low plasma cholesterol concentration correlates with a poor prognosis in septic patients. (**A**) Distribution of total cholesterol levels by 28-day mortality status. Survivors had significantly higher median total cholesterol levels (within 24 hours of ICU admission), p = 0.000518, Wilcoxon rank-sum test. (**B**) Kaplan-Meier survival curves comparing patients with high versus low total cholesterol levels (dichotomized at the median of 133 mg/dL) showed significantly worse 28-day survival in the low-cholesterol group (log-rank p = 0.00035). In the Cox proportional hazards model, adjusted for age, gender, race, SOFA score, and Charlson comorbidity index, higher cholesterol was associated with reduced mortality (adjusted HR = 0.8, 95% CI: 0.67-0.95, p = 0.012).

### Elevation in plasma cholesterol protects against polymicrobial sepsis

To test this hypothesis, we fed C57BL/6J mice with either a high-cholesterol diet (HCD) or a regular diet (RD) for three days. HCD feeding significantly increased plasma cholesterol levels from 78 to 136 mg/dl without altering the cholesterol ester-to-free cholesterol (CE/FC) ratio (**Fig. 2A-D**). Lipoprotein profiling showed increased cholesterol across all lipoprotein fractions in HCD-fed mice (**Fig. 2E-G**). Following cecal ligation and puncture (CLP), HCD-fed mice maintained substantially higher plasma cholesterol concentrations throughout the septic course compared to RD-fed mice (**Fig. 2A-D**). Notably, the CE/FC ratio was increased in the HCD group at 4 hours post-CLP (**Fig. 2D**). Consistent with our clinical observations, HCD feeding significantly increased survival from 52.5% to 90% (**Fig. 2H**).

**Fig. 2.**
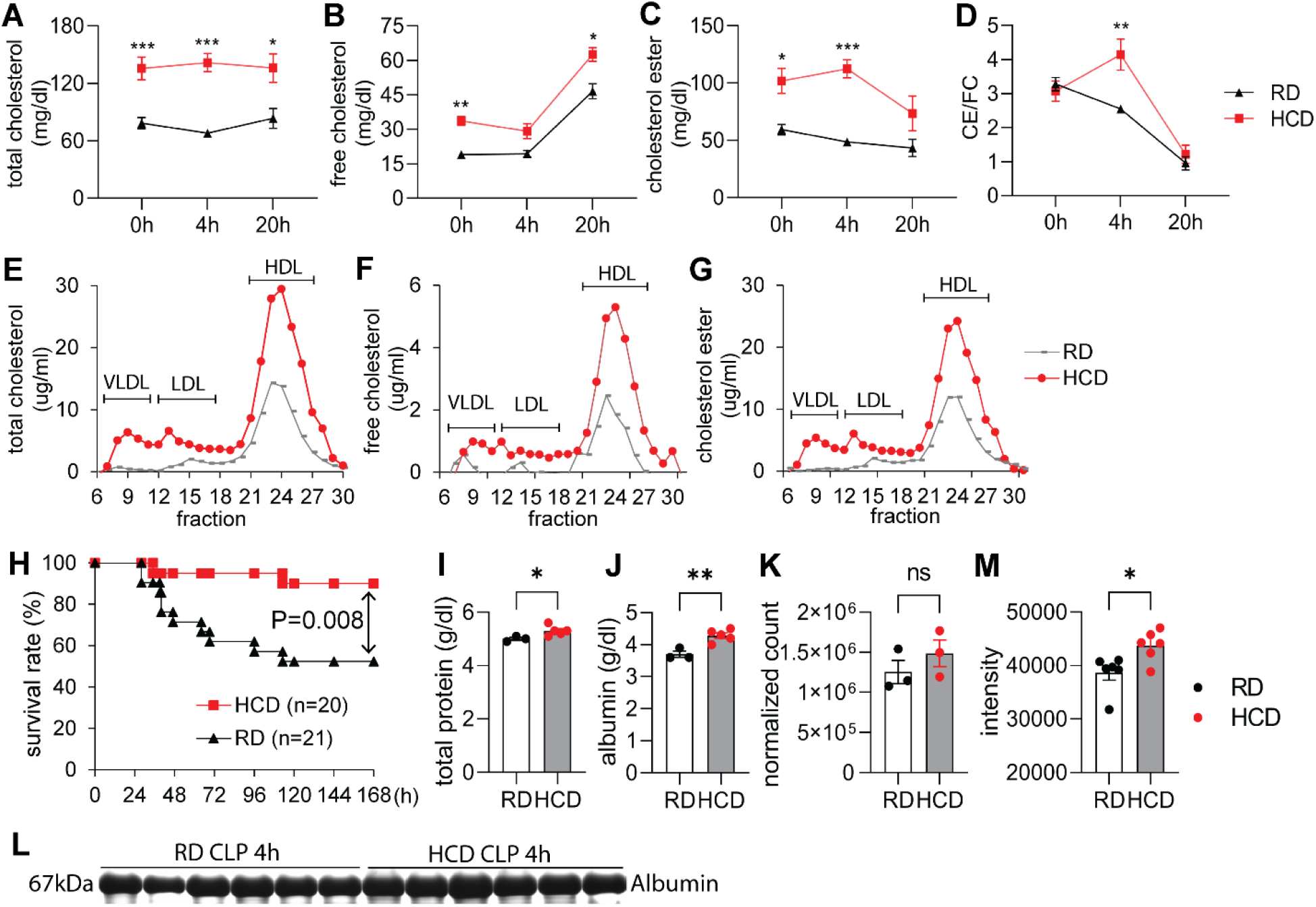
Elevated plasma cholesterol protects against CLP-induced septic death. C57BL/6J mice were fed a regular diet (RD) or high-cholesterol diet (HCD) for 3 days, followed by induction of sepsis via cecal ligation and puncture (CLP). (**A–D**) Serum concentration of (**A**) total cholesterol, (**B**) free cholesterol (FC), (**C**) cholesteryl ester (CE), and (**D**) CE/FC ratio at 0, 4, and 20 hours post-CLP (n = 10–22 per group). (**E–G**) Fast protein liquid chromatography (FPLC) profiles of serum (**E**) total cholesterol, (**F**) free cholesterol (FC), and (**G**) cholesterol ester (CE) levels after 3 days of feeding (n = 2 per group, representative FPLC profiling). (**H**) Survival analysis post-CLP (n = 9–11 per group), assessed by log-rank test. (**I-K**) Serum total protein (**I**) and albumin (**J**) levels, and RNA sequencing analysis of Albumin gene expression in the liver (**K**) following a 3-day diet. (**L-M**) Serum was collected 4 hours after cecal ligation and puncture (CLP), and an equal volume of serum was applied to SDS-PAGE. The gel was stained with Coomassie blue (**L**), and serum albumin was quantified (**M**). Data are presented as mean ± SEM; Statistical significance was determined by two-way ANOVA (**A–D**), Log-Rank test (**H**) or t-test (**I-M**). ^*^p < 0.05, ^**^p < 0.01, ^***^ p < 0.001 for HCD vs. RD comparisons.

To explore protective mechanisms, we assessed organ injury markers—plasma alanine aminotransferase (ALT) for liver injury and blood urea nitrogen (BUN) for kidney damage—and found no significant differences between HCD- and RD-fed mice at 4, 20, and 44 hours post-CLP (**Supplemental Fig. 2A, B**). We next evaluated the inflammatory response by measuring 32 serum cytokines and nitric oxide metabolites (NOx). Only a few cytokines differed between groups (**Supplemental Fig. 2C and Table 2**) and there were no differences in NOx levels (**Supplemental Fig. 2D)**. Flow cytometry analysis revealed minor differences in leukocyte recruitment to the peritoneal cavity (**Supplemental Fig. 3**) and spleen (**Supplemental Fig. 4**). To determine whether these modest changes affected bacterial clearance, we quantified bacterial loads in blood, peritoneal fluid, and organs, and observed no significant differences **(Supplemental Fig. 5**). These findings suggest that elevated plasma cholesterol confers protection through mechanisms other than modulation of inflammation or bacterial clearance.

Because sepsis induces adrenal glucocorticoid (GC) synthesis, which uses cholesterol as a substrate, we measured corticosterone levels. Both groups exhibited a marked rise at 4 hours post-CLP, with no significant differences between HCD and RD mice (**Supplemental Fig. 2E**).

Blood chemistry analysis revealed significantly higher plasma total protein and albumin in HCD-fed mice (**Fig. 2I, J**) without significant change in Albumin gene expression in the liver (**Fig. 2K**). Four hours post-CLP, the HCD-fed mice still had higher albumin levels compared to RD-fed mice (**Fig. 2L, M**). These suggest potential metabolic alterations.

### Elevation in plasma cholesterol induces metabolic reprogramming in the liver that protects against sepsis

To further explore the mechanisms, we performed RNA-seq analysis of liver tissue. As shown in **Fig. 3A**, we identified 558 and 658 unique genes expressed in the RD and HCD groups, respectively. Differential expression analysis revealed 1,471 upregulated and 1,337 downregulated genes in HCD-fed mice compared to RD controls (**Fig. 3B**).

**Fig. 3.**
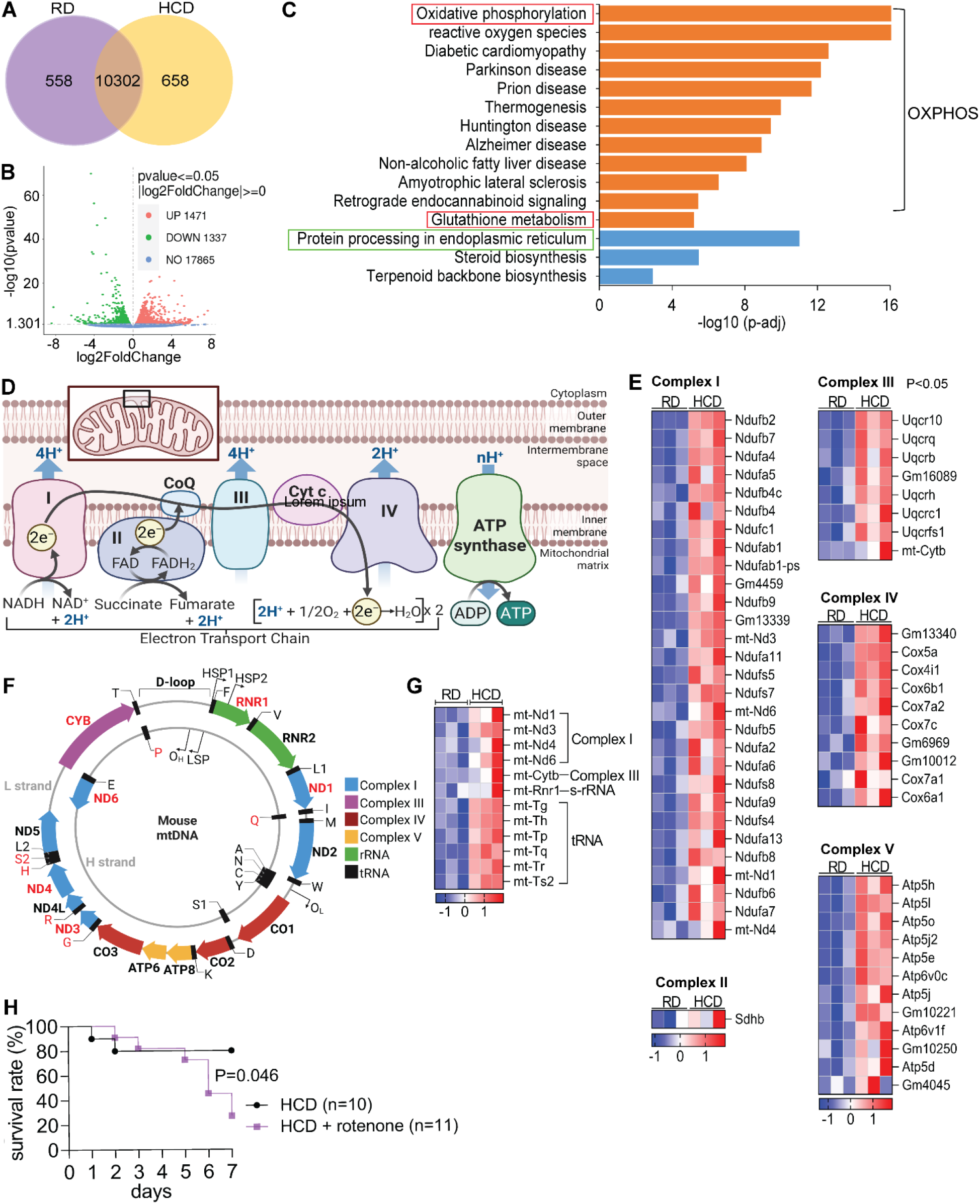
Elevated plasma cholesterol induces metabolic reprogramming in the liver that protects against sepsis. C57BL/6J mice were fed a regular diet (RD) or high-cholesterol diet (HCD) for 3 days, and liver RNA sequencing was conducted. (**A**) Venn diagram showing the distribution of overlapping and unique differentially expressed genes (DEGs) between the HCD and RD groups. (**B**) RNA-seq analysis identifies altered genes in HCD-fed mice. (**C**) Top 12 upregulated pathways identified by KEGG (orange bar) and downregulated pathways (blue bar). (**D**) Schematic of mitochondrial respiratory chain complexes I–V. (**E**) Heatmap of upregulated differentially expressed genes (DEGs) in mitochondrial respiratory chain complexes I–V (p < 0.05). (**F**) Schematic of mitochondrial-encoded genes. (**G**) Heatmap of upregulated DEGs in mitochondrial DNA encoded genes. (**H**) Survival analysis of wild-type mice fed RD or HCD and treated with or without rotenone (a mitochondrial complex I inhibitor) following CLP-induced sepsis. Survival was assessed using the Log-Rank test.

KEGG pathway enrichment analysis highlighted oxidative phosphorylation (OXPHOS) as the most significantly upregulated pathway (adj. P = 5.01 × 10^−17^; **Fig. 3C**). This was accompanied by coordinated transcriptional activation of both nuclear- and mitochondrial-encoded genes across respiratory chain complexes I–V (**Fig. 3D-G**). Of the 65 genes encoding these complexes, 60 were significantly upregulated (**Fig. 3E**), including mitochondrial DNA-encoded genes such as Nd1, Nd3, Nd4, Nd6, and Cytb (**Fig. 3F, G**). Interestingly, the top 11 KEGG pathways —including diabetic cardiomyopathy, prion disease, thermogenesis, Parkinson’s disease, and Alzheimer’s disease—shared substantial overlap in upregulated genes with the OXPHOS pathway (**Supplemental Fig. 6**). This suggests that their enrichment may reflect shared mitochondrial gene activation rather than distinct disease-specific processes. Taken together, these findings indicate that HCD induces a pronounced mitochondrial metabolic shift in the liver, characterized by enhanced OXPHOS activity.

Activation of OXPHOS pathways can increase oxidative stress, prompting us to examine the corresponding antioxidant responses. KEGG pathway analysis identified glutathione metabolism as the most significantly upregulated pathway after OXPHOS (adj. P = 2.32 × 10^−7^; **Fig. 3C**). A broad array of antioxidant defense genes was upregulated (**Fig. 4A, B**). Key enzymes involved in glutathione biosynthesis and recycling were upregulated, including Gclc and Gclm (which form the rate-limiting enzyme glutamate-cysteine ligase), and Gss (glutathione synthetase). Glutathione peroxidases (Gpx1, Gpx4, Gpx4-ps2) and Prdx6 were also elevated, indicating enhanced enzymatic reduction of hydrogen peroxide and lipid peroxides. Multiple glutathione S-transferases (Gstm2-ps1, Gsta2, Gsta4, Gstm1, Mgst3, Gsto1, Gstt2, Gstm3, Gstt3, Gstp3, Gsta1, Gstt1, Gstk1), catalyze the conjugation of glutathione to electrophilic compounds, facilitating their detoxification. In addition, elevated expression of Idh2 and Ggt6 further supports increased NADPH production and glutathione turnover, essential for maintaining the reduced state of glutathione. Together, these changes reflect a coordinated antioxidant response aimed at maintaining redox balance under elevated mitochondrial activity.

**Fig. 4.**
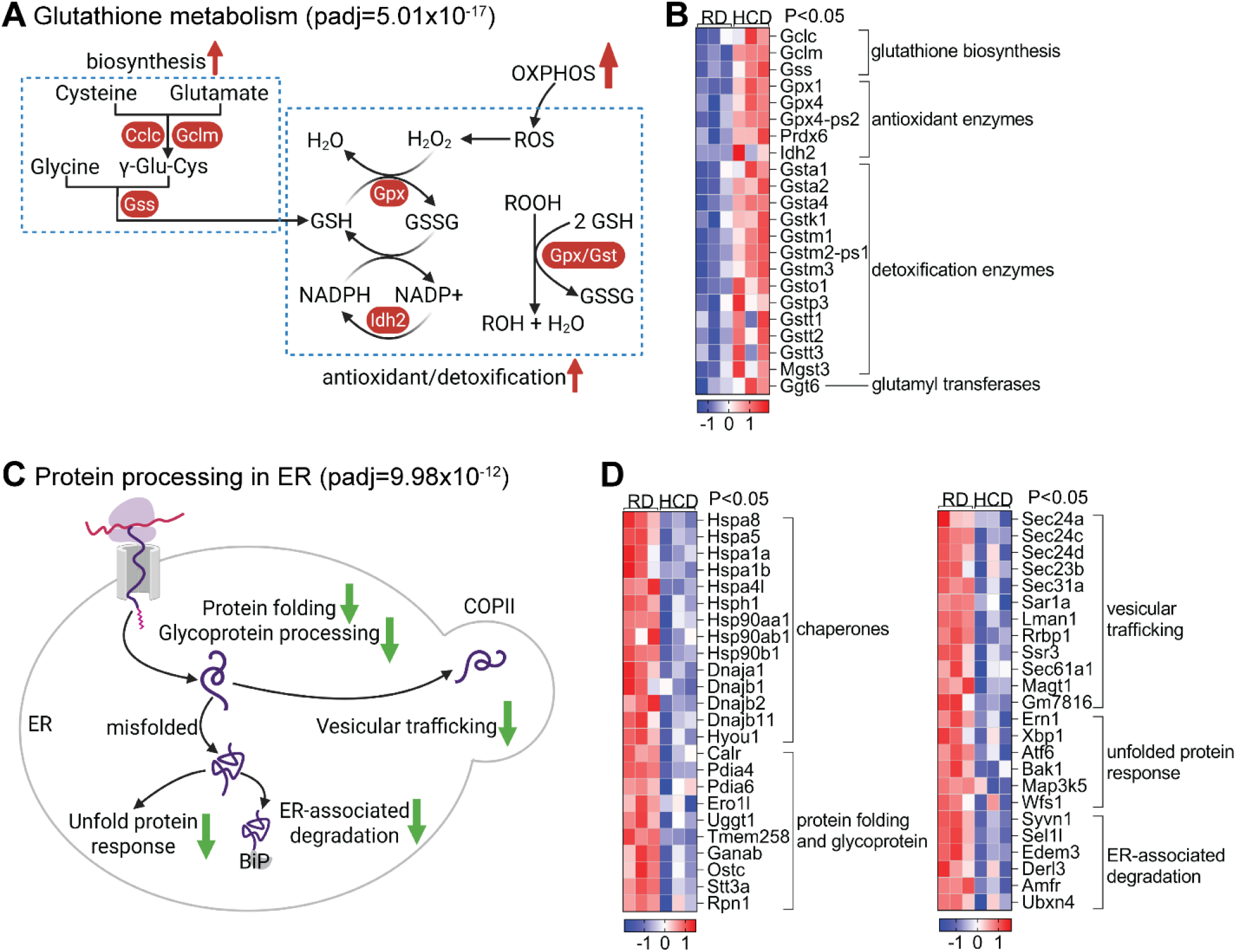
Elevated plasma cholesterol promotes metabolic reprogramming - upregulating glutathione metabolism and downregulating protein processing in endoplasmic reticulum (ER). RNA sequencing analysis was conducted on liver tissues from mice fed a high-cholesterol diet (HCD) or a regular diet (RD). (**A**) KEGG pathway enrichment analysis revealed significant upregulation of the glutathione metabolic pathway. (**B**) Heatmap showing upregulated differentially expressed genes (DEGs) in glutathione biosynthesis and in antioxidation (p < 0.05). (**C**) KEGG pathway enrichment analysis revealed significant downregulation of protein processing in ER pathway. (**D**) Heatmap showing downregulated DEGs in the ER protein processing pathway. (p < 0.05).

In contrast to the upregulation of oxidative metabolism, KEGG analysis identified protein processing in the endoplasmic reticulum (ER) as the most significantly downregulated pathway in HCD-fed mice (adj. P = 3.16 × 10^−14^; **Fig. 3C**). This suppression was characterized by broad transcriptional downregulation of genes involved in ER function, protein folding and degradation, suggesting a metabolic shift away from proteostasis toward energy conservation (**Fig. 4C, D)**. Key molecular chaperones—including Hspa5, Hspa8, Hspa1a, Hspa1b, Hspa4l, Hsp90aa1, Hsp90ab1, Hsp90b1, Hsph1, Dnaja1, Dnajb1, Dnajb2, and Dnajb11—were significantly suppressed, indicating reduced protein folding capacity. Similarly, unfolded protein response (UPR) regulators such as Xbp1, Atf6, Ern1, Hyou1, and Map3k5 were downregulated, reflecting diminished ER stress signaling. Components of the ER-associated degradation (ERAD) pathway—including Syvn1, Sel1l, Derl3, Amfr, and Ubxn4—were also reduced, along with genes involved in vesicular trafficking (Sec24a, Sec24c, Sec24d, Sec23b, Sec31a, Sar1a, Sec61a1, Ssr3), suggesting a global decline in secretory activity. Additionally, genes responsible for oxidative protein folding and glycoprotein processing (Pdia4, Pdia6, Ero1l, Uggt1, Ostc, Stt3a, Ganab, Rpn1, Lman1, Tmem258, Ggt6, Wfs1) were consistently downregulated. Functionally, this coordinated suppression of ER activity may reduce energy expenditure associated with protein degradation, thereby preserving essential plasma proteins during systemic stress. Supporting this, plasma total protein and albumin levels were significantly higher in HCD-fed mice compared to RD controls, suggesting improved protein retention during sepsis.

Finally, to assess functional relevance, we treated HCD-fed mice with rotenone, a mitochondrial complex I inhibitor, and evaluated survival following CLP-induced sepsis. Rotenone treatment significantly reduced survival compared to untreated controls, confirming that mitochondrial respiration is a critical mediator of cholesterol-induced hepatic reprogramming and host defense (**Fig. 3H**).

Collectively, these findings demonstrate that elevated plasma cholesterol drives a comprehensive hepatic metabolic reprogramming—characterized by activation of mitochondrial respiration and antioxidant defenses, alongside suppression of ER proteostasis. This coordinated shift prioritizes energy efficiency and redox balance, thereby enhancing systemic resilience to sepsis-induced stress (**Fig. 5**).

**Fig. 5.**
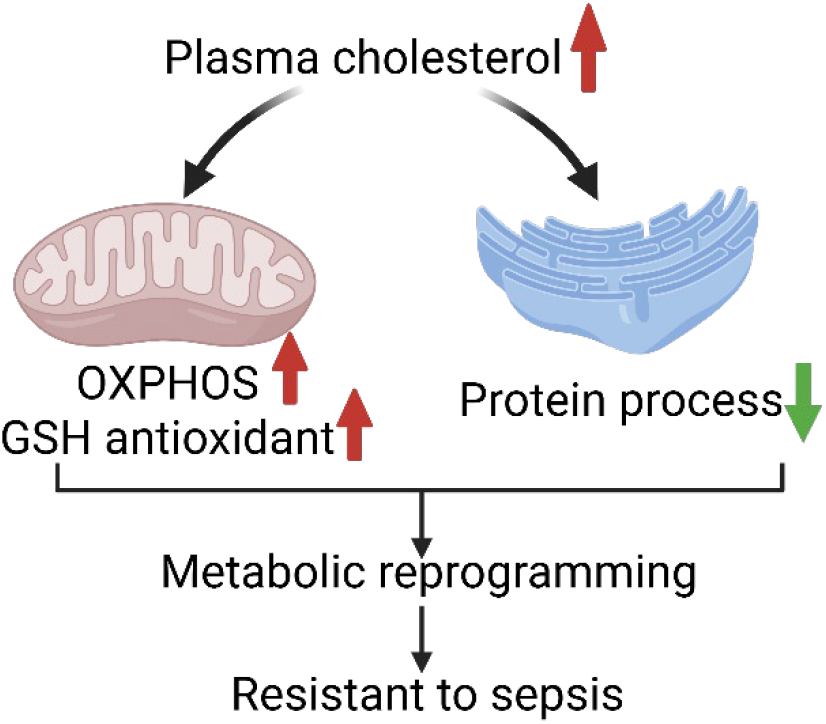
Schematic model of cholesterol-mediated protection in sepsis. Elevated plasma cholesterol confers a survival advantage in sepsis by promoting hepatic metabolic reprogramming. This includes upregulation of mitochondrial oxidative phosphorylation and glutathione-mediated antioxidant defenses, alongside attenuation of the energetic burden imposed by endoplasmic reticulum (ER) proteostasis. These coordinated adaptations enhance cellular energy balance and reduce oxidative stress, contributing to improved outcomes in sepsis. Notably, this protective effect occurs largely independent of classical immune pathways, highlighting hepatic bioenergetics as a novel and potentially targetable mechanism for sepsis therapy. Created with Biorender.com.

## Discussion

The longstanding association between low plasma cholesterol and poor outcomes in sepsis has led to speculation about a protective role for cholesterol during systemic infection (5, 7). However, previous clinical studies were limited by small sample sizes and inadequate adjustment for confounding variables (14-16). By leveraging a large, well-curated cohort from the MIMIC-IV database, our study provides strong support for this clinical association and provides mechanistic insights with complementary experimental data.

Our analysis confirms that hypocholesterolemia is independently associated with increased mortality in critically ill septic patients. Specifically, non-survivors had significantly lower plasma total cholesterol compared to survivors, and patients in the low-cholesterol group exhibited worse 28-day ICU survival after adjusting for age, gender, race, SOFA score within the first 24 hours, and Charlson comorbidity index. The relationship persists across different SOFA score ranges, suggesting it’s not simply explained by disease severity.

In a murine model of sepsis, dietary cholesterol supplementation elevated plasma cholesterol levels and significantly improved survival, independent of classical immune responses such as cytokine production, leukocyte recruitment, or bacterial clearance. This observation shifts the focus from immunomodulation to metabolic adaptation as a critical determinant of host resilience. Mechanistically, cholesterol induced a hepatic metabolic reprogramming characterized by enhanced oxidative phosphorylation and antioxidant defenses, coupled with suppression of endoplasmic reticulum (ER) proteostasis. These coordinated changes optimize energy production, mitigate oxidative stress, and reduce protein wasting during sepsis. The functional relevance of this metabolic shift was confirmed by pharmacological inhibition of mitochondrial respiration, which abolished the survival benefit conferred by cholesterol supplementation, underscoring the central role of mitochondrial metabolism in mediating host protection.

Sepsis is known to induce a metabolic shift from mitochondrial OXPHOS to glycolysis, particularly in immune and parenchymal cells. While glycolysis supports rapid ATP generation and acute immune activation, it is energetically inefficient and contributes to immunosuppression and organ dysfunction over time. Restoring mitochondrial OXPHOS has thus emerged as a promising therapeutic strategy. Our findings reinforce this concept. Although cholesterol was used as a pre-treatment in our study, the transcriptional programs identified—especially those enhancing mitochondrial function and antioxidant capacity—represent potential therapeutic targets. Modulating these pathways through pharmacological agents or tailored nutritional interventions may improve host resilience, even in the absence of prior cholesterol elevation. Supporting this concept, early preclinical studies have shown that pharmacologic activation of the pyruvate dehydrogenase complex (PDC) using dichloroacetate (DCA) reactivates mitochondrial respiration, improves energy metabolism in hepatocytes and splenocytes, and significantly enhances survival in sepsis models (17).

Notably, our findings contrast with several prior studies reporting that high-fat or high-cholesterol diets impair mitochondrial OXPHOS (18-21). Several factors may explain this discrepancy. First, most previous studies employed long-term dietary interventions spanning several months, whereas our study utilized a short-term feeding protocol. Second, our HCD included cholate, a bile acid that markedly elevates plasma cholesterol levels and may amplify metabolic effects. Third, earlier studies relied on targeted approaches such as RT-PCR or Western blotting, which focus on a limited set of mitochondrial markers and may miss broader transcriptional changes. Moreover, RT-PCR normalization typically assumes stable expression of housekeeping genes, yet our RNA-seq data revealed that β-actin was significantly upregulated (1.5-fold) in HCD-fed mice compared to RD controls. In contrast, our use of unbiased RNA sequencing enabled comprehensive profiling of transcriptional changes, revealing systemic metabolic reprogramming—including robust upregulation of OXPHOS genes across all mitochondrial complexes and antioxidative pathways, alongside suppression of ER proteostasis pathways—that targeted methods may overlook.

Our study has several limitations. First, the observational nature of the clinical data precludes causal inference, and residual confounding cannot be fully excluded. The use of cholesterol-lowering medication prior to ICU admission was unknown and thus not considered in the analysis. Second, the murine model may not fully recapitulate the complexity and heterogeneity of human sepsis, particularly with respect to comorbidities, microbiota, and immune responses. Finally, while our transcriptomic data provide valuable insights into metabolic reprogramming, further studies are needed to validate these findings at the protein and functional levels, and to determine whether similar pathways are activated in human tissues during sepsis.

In summary, our study identifies cholesterol as a critical regulator of host resilience during sepsis. By promoting hepatic metabolic reprogramming—enhancing mitochondrial respiration and antioxidant defenses, while alleviating the energetic burden of ER proteostasis—cholesterol supplementation confers a survival advantage that appears independent of classical immune pathways. These findings offer mechanistic insight into longstanding clinical observations linking hypocholesterolemia with poor sepsis outcomes and suggest that targeting hepatic metabolism may represent a novel therapeutic strategy. Future studies should explore pharmacological approaches that mimic the beneficial metabolic programs induced by cholesterol, potentially offering new avenues to improve outcomes in sepsis and related critical illnesses.

## Acknowledgments

Q Wang and X-A Li designed the experiments, Q Wang, L. Guo, J Xue, D Hao, M Ito, performed experiments and data analysis; C Wu conducted the retrospective study with the help from Q Wang and R Reese; B Huang verified statistical analysis; Q Wang verified the data and wrote the original manuscript; X-A Li acquired funding and edited the manuscript. C Wu edited the manuscript. All authors have read and agreed to submit the manuscript. We thank Stacey A. Slone for statistical support (Data Analytics Core of the PADS Hub at the University of Kentucky).

## Funding and additional information

This study was supported by Grants NIH R35GM141478 and VA I01BX006408 (to X-A Li), and supported by the Shared Resource(s) of the University of Kentucky Markey Cancer Center (P30CA177558). C.W. is supported by NIH R01HL177556. Its contents are solely the responsibility of the authors and do not necessarily represent the official views of the National Institutes of Health or VA.

## Abbreviations

ALT: alanine aminotransferase
ATP: adenosine triphosphate
BUN: blood urea nitrogen
CE: cholesteryl ester
CLP: cecal ligation and puncture
DEG: differentially expressed gene
ER: endoplasmic reticulum
ERAD: ER-associated degradation
FC: free cholesterol
FPLC: fast protein liquid chromatography
GC: glucocorticoid
Gclc: glutamate-cysteine ligase catalytic subunit
Gclm: glutamate-cysteine ligase modifier subunit
Gpx: glutathione peroxidase
GR: glucocorticoid receptor
GSH: glutathione (reduced form)
GSSG: glutathione disulfide (oxidized form)
HCD: high-cholesterol diet
HR: hazard ratio
HSP: heat shock protein
ICU: intensive care unit
KEGG: Kyoto Encyclopedia of Genes and Genomes
MAPK: mitogen-activated protein kinase
MIMIC-IV: Medical Information Mart for Intensive Care, version IV
mRNA: messenger ribonucleic acid
NADPH: nicotinamide adenine dinucleotide phosphate
NOx: nitrate oxide
OXPHOS: oxidative phosphorylation
PBS: phosphate-buffered saline
PDC: pyruvate dehydrogenase complex
PMN: polymorphonuclear neutrophil
qPCR: quantitative polymerase chain reaction
RD: regular diet
RNA-seq: RNA sequencing
ROS: reactive oxygen species
SEM: standard error of the mean
SQL: structured query language
TC: total cholesterol
UPR: unfolded protein response

