## Supplemental for "Elevated plasma cholesterol causally improves sepsis outcome by promoting hepatic metabolic reprogramming"

**Supplemental Tables:**

**Supplemental Table 1.** Baseline characteristics of patients

| Characteristic | Overall N = 2,787[1] | Survivor N = 2,211[1] | Nonsurvivor N = 576[1] | p-value[2] |
|---|---|---|---|---|
| **Age (years)** | 66.1 (15.3) | 64.9 (15.3) | 70.6 (14.3) | <0.001 |
| **Gender** | | | | 0.6 |
| Female | 1,129 (41%) | 889 (40%) | 240 (42%) | |
| Male | 1,658 (59%) | 1,322 (60%) | 336 (58%) | |
| **SOFA Score** | 5.5 (3.1) | 5.4 (3.1) | 6.1 (3.4) | <0.001 |
| **Charlson Comorbidity Index** | 5.8 (2.9) | 5.5 (2.8) | 6.8 (2.9) | <0.001 |
| **Total Cholesterol (mg/dL)** | 138.8 (54.5) | 140.2 (53.8) | 133.5 (56.5) | 0.010 |
| **HDL Cholesterol (mg/dL)** | 39.4 (17.8) | 39.3 (17.7) | 39.8 (18.3) | 0.5 |
| **LDL Cholesterol (mg/dL)** | 75.5 (40.1) | 76.1 (39.7) | 73.2 (41.6) | 0.2 |

[1] Mean (SD); n (%)

[2] Welch Two Sample t-test; Pearson's Chi-squared test

SOFA, Sequential Organ Failure Assessment; HDL, High-Density Lipoprotein; LDL, Low-Density Lipoprotein

**Supplemental Table 2.** Cytokine levels of B6 mice with or without HCD before CLP and at 4, 20, and 44hours after CLP-induced sepsis.

| | no CLP | | CLP 4h | | CLP 20h | | CLP 44h | | P-values | | | |
|---|---|---|---|---|---|---|---|---|---|---|---|---|
| Cytokines | RD | HCD | RD | HCD | RD | HCD | RD | HCD | no CLP | 4h | 20h | 44h |
| Eotaxin | 408.63±63.5 | 438.12±23.25 | 1843.62±132.02 | 1460.06±268.08 | 645.39±95.74 | 1018.02±170.91 | 442.62±28.44 | 460.42±31.58 | 0.415 | 0.224 | 0.081 | 0.68 |
| G-CSF | 289.78±47.24 | 259.49±21.91 | 58728.65±1722.13 | 59299.16±2061.77 | 40750.94±1042.64 | 39788.4±1288.4 | 40068.29±812.39 | 37106.52±735.43 | 0.282 | 0.834 | 0.57 | 0.014* |
| GM-CSF | 10.24±2.17 | 10.5±2.57 | 153.19±13.47 | 138.99±8.61 | 65.4±9.99 | 55.81±6.19 | 51.08±7.52 | 55.41±12.1 | 0.877 | 0.388 | 0.431 | 0.765 |
| IFNy | 0.47±0.11 | 0.57±0.33 | 50.13±18.68 | 33.59±8.74 | 17.9±9.4 | 10.82±2.7 | 20.94±10 | 17.13±7.91 | 0.593 | 0.437 | 0.489 | 0.768 |
| IL-1a | 69.78±14.51 | 95.46±26.62 | 682.73±37.62 | 790.51±115.57 | 903.83±442.1 | 471.45±71.83 | 369.39±50.35 | 478.37±110.33 | 0.13 | 0.397 | 0.365 | 0.384 |
| IL-1B | 3.42±0.7 | 3.66±0.66 | 87.81±18.05 | 38.26±6.19 | 49.6±33.96 | 14.22±4.36 | 11.63±1.66 | 11.99±1.63 | 0.635 | 0.025* | 0.335 | 0.879 |
| IL-2 | 7.83±3.17 | 7.2±1.41 | 94.59±30.28 | 68.47±22.61 | 88.76±47.6 | 26.94±4.21 | 42.04±22.64 | 46.69±26.55 | 0.727 | 0.499 | 0.236 | 0.895 |
| IL-3 | 1.1±0.32 | 0.97±0.09 | 5.75±1.63 | 7.07±1.74 | 12.58±9.46 | 2.04±0.8 | 3.73±1.67 | 10.12±5.55 | 0.452 | 0.587 | 0.303 | 0.292 |
| IL-4 | 0.24±0.05 | 0.28±0.09 | 4.95±1.59 | 4.41±1.27 | 2.1±0.61 | 1.71±0.23 | 3.07±1.55 | 1.59±0.34 | 0.409 | 0.794 | 0.567 | 0.372 |
| IL-5 | 6.38±1.69 | 3.29±0.69 | 632.99±91.1 | 486.31±75.8 | 54.36±11.1 | 92.53±31.94 | 25.82±6.89 | 17.43±3.7 | 0.013* | 0.233 | 0.286 | 0.302 |
| IL-6 | 2.23±0.22 | 2.81±1.4 | 34267.04±4316.09 | 29306.14±5130.42 | 2798.2±1366.91 | 4671.77±2756.19 | 876.31±180.25 | 1061.75±329.32 | 0.443 | 0.47 | 0.554 | 0.628 |
| IL-7 | 2.52±0.62 | 2.74±0.89 | 11.5±2.61 | 13.03±2.29 | 31.75±25.05 | 6.26±2.29 | 4.21±1.88 | 35.94±27.76 | 0.697 | 0.666 | 0.344 | 0.28 |
| IL-9 | 6.32±9.4 | 1.83±0.29 | 239.89±58.49 | 408.22±182.27 | 153.14±42.35 | 48.59±14.56 | 41.44±13.77 | 56.4±12.11 | 0.383 | 0.401 | 0.046* | 0.426 |
| IL-10 | 1.59±0.37 | 1.45±0.63 | 3585.11±1171.87 | 1006.97±263.36 | 415.45±226.36 | 1492.72±1375.19 | 47.97±19.78 | 36.98±19.95 | 0.723 | 0.058 | 0.461 | 0.7 |
| IL-12 (p40) | 46.63±63.85 | 15.87±4.11 | 311.38±269.51 | 90.67±71.29 | 202.88±189.33 | 18.99±10.59 | 1147.54±1125.42 | 56.96±38.31 | 0.38 | 0.446 | 0.364 | 0.358 |
| IL-12 (p70) | 13.7±2.12 | 16.16±2.56 | 340.82±55.67 | 299.73±65.36 | 113.11±34.39 | 82.28±15.05 | 109.56±56.99 | 51.18±10.33 | 0.17 | 0.639 | 0.431 | 0.338 |
| IL-13 | 9.47±1.46 | 10.27±2.25 | 270.37±26.28 | 220.16±23.51 | 66.39±11.57 | 68.32±13.36 | 44.91±6.03 | 53.7±11.09 | 0.566 | 0.173 | 0.915 | 0.496 |
| IL-15 | 17.82±5.36 | 22.05±9.37 | 183.06±20.49 | 197.65±29.6 | 339.59±271.08 | 124.68±22.08 | 84.94±18.78 | 289.75±238.59 | 0.456 | 0.691 | 0.455 | 0.412 |
| IL-17 | 1.74±1.3 | 0.67±0.25 | 159.76±34.83 | 109.68±15.35 | 20.27±8.24 | 39.54±21.85 | 13.72±5.3 | 10.37±1.77 | 0.161 | 0.212 | 0.428 | 0.562 |
| IP-10 | 54.52±6.5 | 59.22±3.84 | 995.95±224.16 | 485.64±66.94 | 159.75±29.11 | 180.07±47.26 | 101.51±5.28 | 112.51±10.2 | 0.247 | 0.053 | 0.72 | 0.354 |
| KC | 232.49±29.18 | 283.29±56.05 | 35275.7±2604.38 | 19801.78±3847.76 | 10601.82±2200.51 | 9937.33±2615.7 | 2163.46±738.8 | 1411.42±299.12 | 0.149 | 0.005* | 0.849 | 0.364 |
| LIF | 0.14±0.22 | 0.06±0.03 | 20±2.67 | 14.42±1.75 | 9.63±4.42 | 10.95±4.38 | 2.67±0.87 | 6.72±4.96 | 0.51 | 0.1 | 0.835 | 0.439 |
| LIX | 366.03±87 | 601.55±367.04 | 7765.23±1732.49 | 5961.92±1776.38 | 4242.47±1504.19 | 4419.79±1364.78 | 1694.03±600.36 | 2475.43±758.75 | 0.262 | 0.477 | 0.932 | 0.43 |
| MCP-1 | 35.03±12.61 | 30.69±5.63 | 3381.41±890.57 | 2164.75±465.35 | 629.61±258.33 | 597.9±174.51 | 128.84±20.82 | 95.34±10.29 | 0.55 | 0.247 | 0.921 | 0.172 |
| M-CSF | 8.09±4.55 | 4.41±2.27 | 10.41±1.39 | 38.34±30.23 | 69.61±57.71 | 7.06±1.44 | 20.12±8.21 | 43.31±27.15 | 0.189 | 0.383 | 0.314 | 0.43 |
| MIG | 64.57±5.18 | 64.83±4.28 | 243±56.85 | 165.54±38.15 | 94.98±26.48 | 173.46±53.35 | 20.56±3.31 | 30.16±5.3 | 0.941 | 0.275 | 0.213 | 0.144 |
| MIP-1a | 47.21±7.71 | 48.23±3.35 | 1592.02±319.87 | 1017.02±149.84 | 422.23±64.34 | 782.74±261.31 | 377.04±94.55 | 208.72±49.7 | 0.8 | 0.128 | 0.213 | 0.138 |
| MIP-1B | 41.14±4.18 | 46.03±4.28 | 6092.77±1913.31 | 2854.11±627.46 | 378.55±115.99 | 879.43±646.69 | 253.97±86.72 | 148.74±43.06 | 0.133 | 0.136 | 0.466 | 0.296 |
| MIP-2 | 64.76±13.28 | 80.71±20.89 | 7370.7±2195.21 | 1691.72±392.68 | 442.48±127.91 | 2698.97±2445.57 | 123.43±63.33 | 333.74±170.06 | 0.231 | 0.03* | 0.384 | 0.268 |
| RANTES | 21.52±5.72 | 18.31±6.28 | 125.66±13.85 | 118.14±15.64 | 89.38±39.4 | 57.57±13.83 | 50.02±12.6 | 62.74±17.99 | 0.468 | 0.723 | 0.466 | 0.57 |
| TNFa | 5.77±1.19 | 5.78±1.37 | 160.21±25.47 | 121.04±14.91 | 83.9±28.03 | 68.34±12.97 | 36.99±3.74 | 33.19±3.23 | 0.989 | 0.205 | 0.625 | 0.452 |
| VEGF | 0.41±0.05 | 0.36±0.02 | 5.15±1.82 | 13.79±7.61 | 3.09±1.04 | 1.17±0.11 | 1.23±0.15 | 1.5±0.23 | 0.149 | 0.298 | 0.108 | 0.355 |

**Supplemental Table 2.** Cytokine levels of B6 mice with or without HCD before CLP and at 4, 20, and 44hours after CLP-induced sepsis. Data are presented as mean ± SEM (n=6-11 for each group). *$p<0.05$ compared between the HCD and RD groups, assessed by Student's t-test.

**Supplemental Figures:**

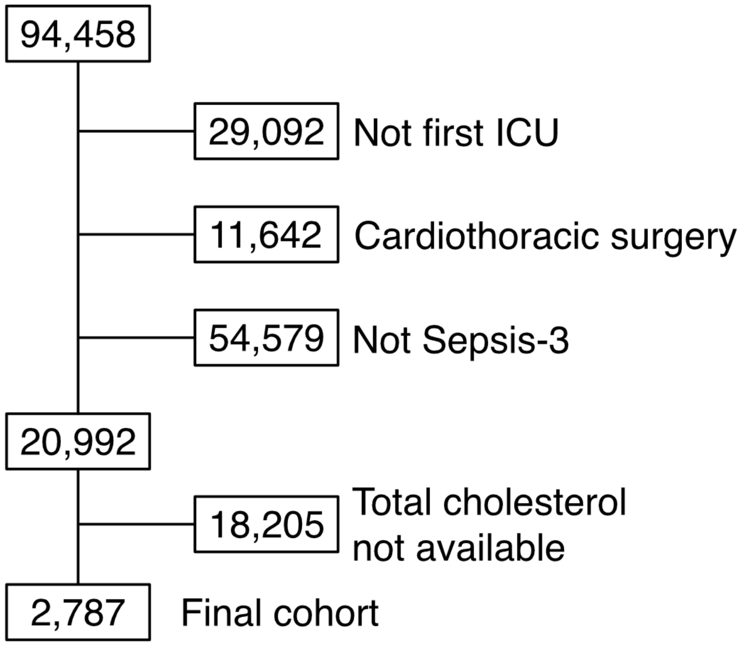


**Supplemental Fig. 1.** Flowchart of study patients

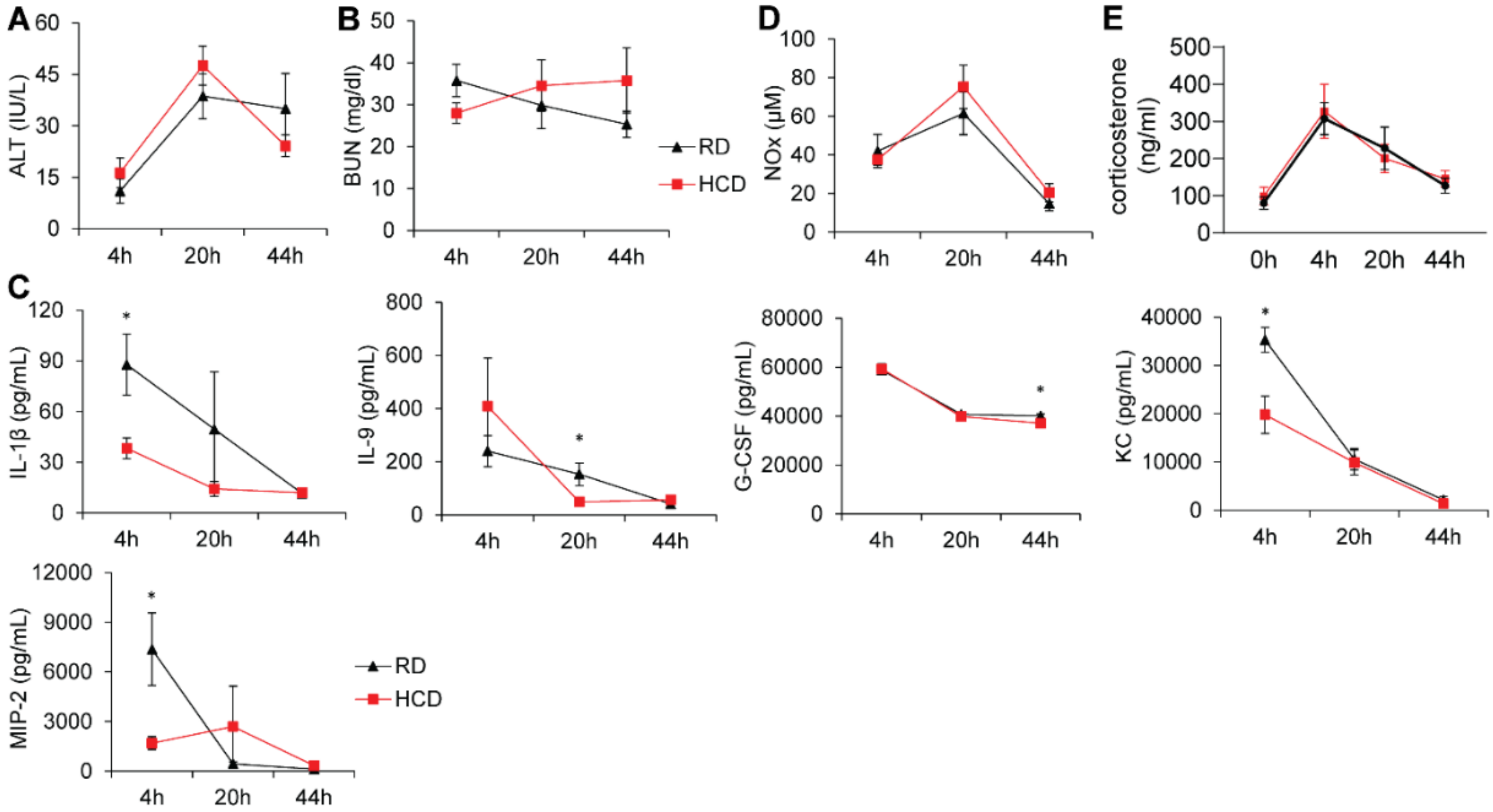


**Supplemental Fig. 2. High-cholesterol diet minimally affects inflammatory response in sepsis.** C57BL/6J mice were fed a regular diet (RD) or high-cholesterol diet (HCD) for 3 days and underwent cecal ligation and puncture (CLP) to induce sepsis. (**A-B**) Serum alanine aminotransferase (ALT) and blood urea nitrogen (BUN) levels post-CLP (n = 9–11/group). (**C**) Serum cytokine levels of IL-1β, IL-9, G-CSF, KC, and MIP-2 in RD- and HCD-fed mice at 4, 20, and 44 hours post-CLP. (**D–E**) Serum levels of (**D**) nitrate/nitrite (NOx), (**E**) corticosterone at the indicated timepoints (n = 4–11 per group). Data are mean ± SEM. Statistical significance was determined by two-way ANOVA. *$p < 0.05$ for HCD vs. RD comparisons.

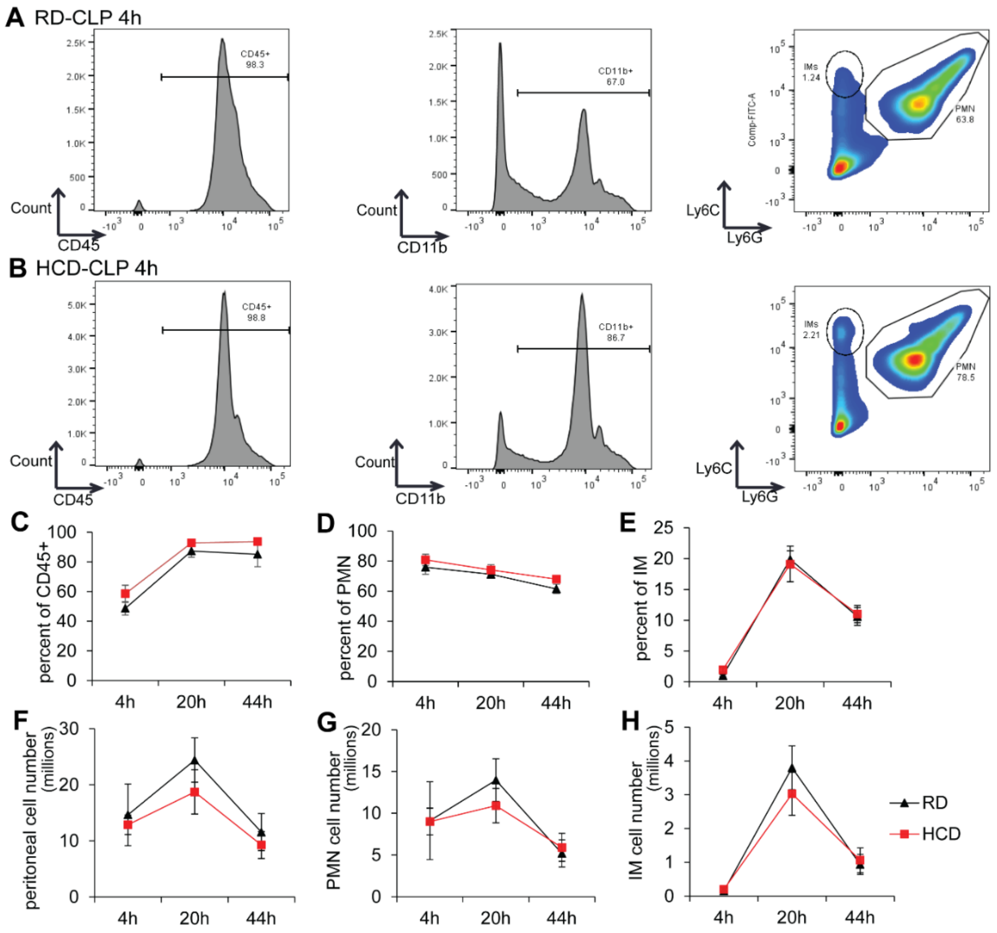


**Supplemental Fig. 3. High-cholesterol diet (HCD) rarely affects CLP-induced leukocyte recruitment to the peritoneal cavity.** C57BL/6J mice were fed a regular diet (RD) or high-cholesterol diet (HCD) for 3 days, followed by induction of sepsis via cecal ligation and puncture (CLP). Peritoneal fluid was collected from mice at 4, 20, and 44 hours post-CLP. (**A, B**) Flow cytometry analysis of peritoneal leukocytes from HCD-fed (**A**) and regular diet (RD)-fed (**B**) mice. Peritoneal neutrophils (PMNs, CD45+CD11b+ Ly6C$^{hi}$ Ly6G+) and inflammatory monocytes (IMs, CD45+CD11b+ Ly6C$^{hi}$ Ly6G-) were gated by Ly6G and Ly6C expression on CD45+ cells. (**C–E**) Percentages of CD45$^{+}$ cells (**C**), PMNs (**D**), and IMs (**E**). (**F**) Peritoneal cell counts. (**G, H**) Numbers of recruited PMNs (**G**) and IMs (**H**). Data are presented as mean ± SEM. Statistical analysis was performed using two-way ANOVA.

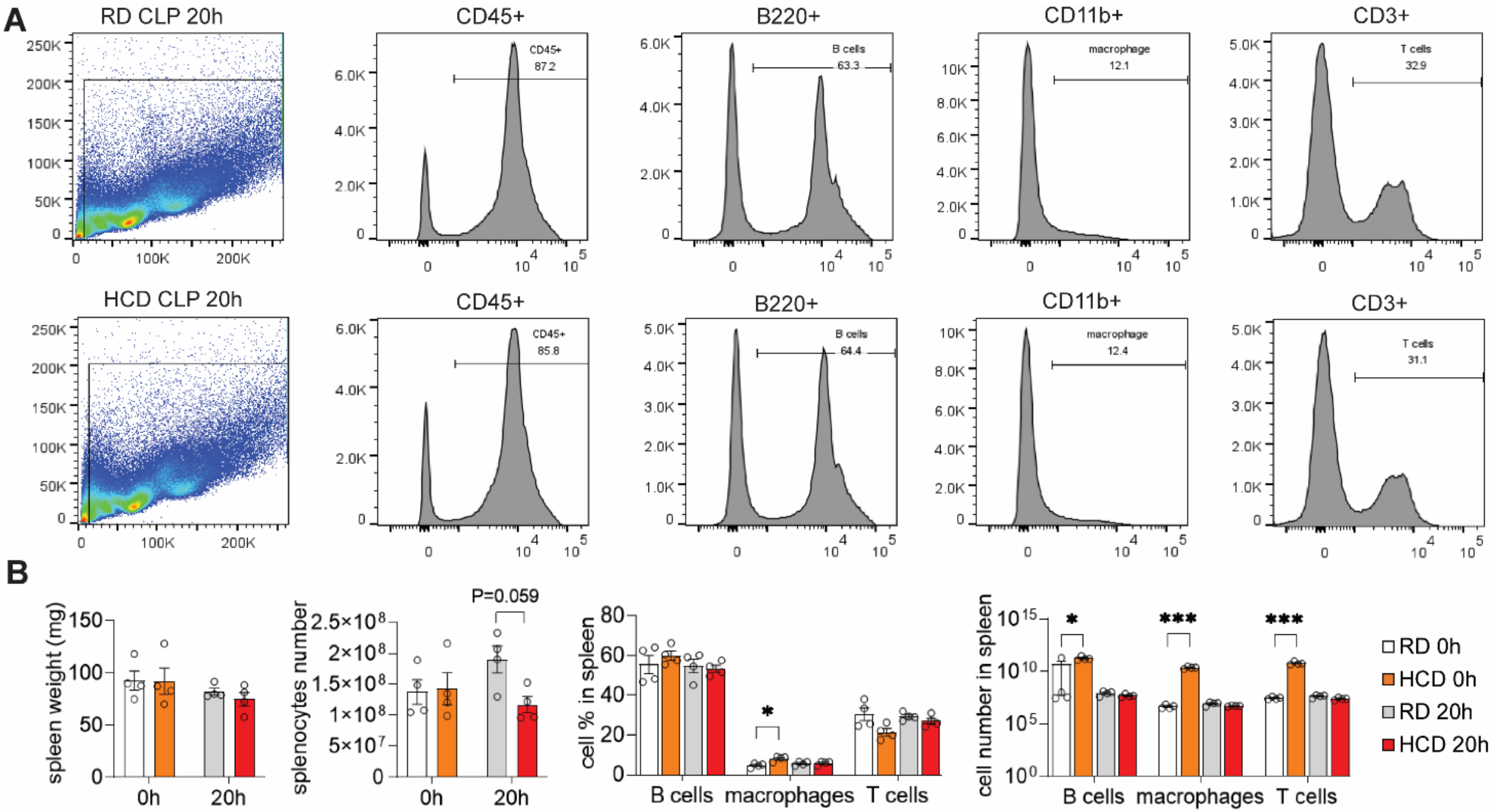


**Supplemental Fig. 4. High-cholesterol diet moderately increases immune cell numbers in the spleen.** C57BL/6J mice were fed a regular diet (RD) or high-cholesterol diet (HCD) for 3 days, followed by induction of sepsis via cecal ligation and puncture (CLP). Spleens were harvested for flow cytometric analysis. (**A**) Representative flow cytometry gating strategy for splenocytes at 20 hours post-CLP, identifying B cells (B220+), macrophages (CD11b+), and T cells (CD3+). (**B**) Quantification of splenic T cells, B cells, and macrophages in RD- and HCD-fed mice at the indicated timepoints (n = 4 per group). Data are mean ± SEM. Statistical significance was determined by two-way ANOVA. *$p < 0.05$, *** $p < 0.001$ for HCD vs. RD comparisons.

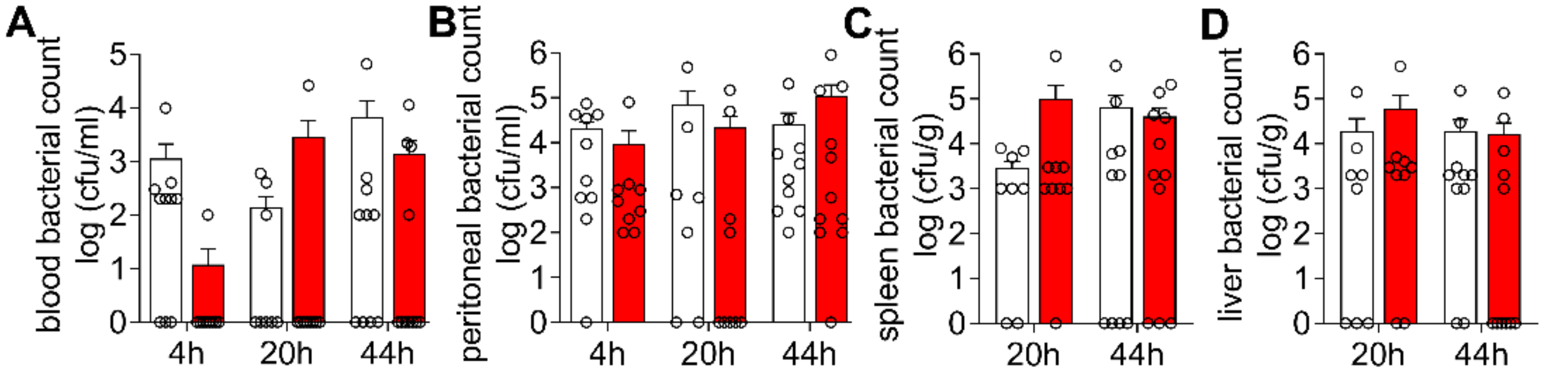


**Supplemental Fig. 5. High cholesterol diet does not alter bacterial loads in blood and organs during sepsis.** C57BL/6J mice were fed a regular diet (RD) or high-cholesterol diet (HCD) for 3 days, followed by induction of sepsis via cecal ligation and puncture (CLP). Bacterial loads were measured in (**A**) blood, (**B**) peritoneal fluid, (**C**) spleen, and (**D**) liver at indicated time points post-CLP (n = 9–11/group). Data are presented as mean ± SEM; Statistical analyses were performed using two-way ANOVA; no significant differences were observed between HCD and RD groups.

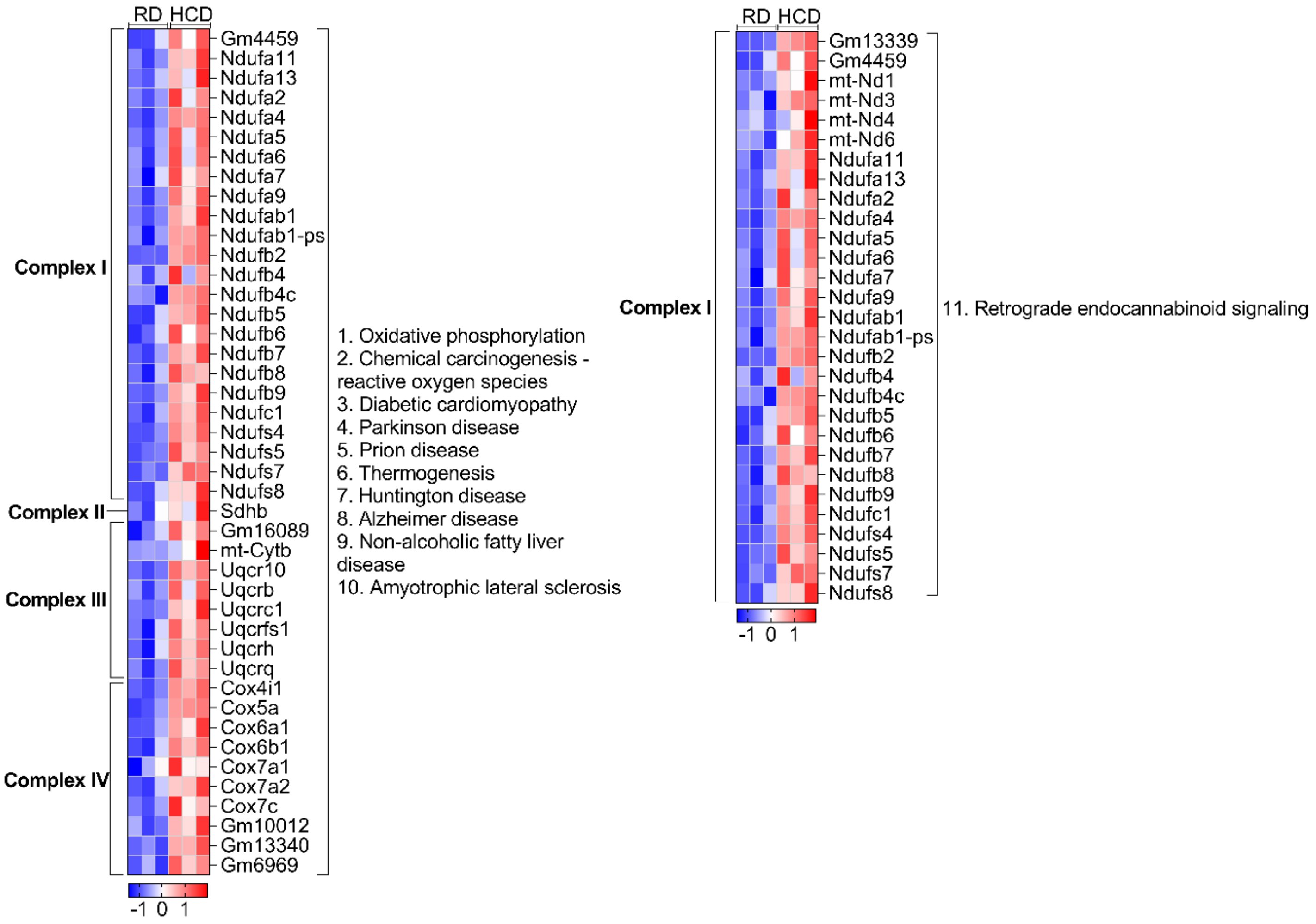


**Supplemental Fig. 6. Differentially expressed genes (DEGs) shared by the top 11 KEGG pathways.** C57BL/6J mice were fed a regular diet (RD) or high-cholesterol diet (HCD) for 3 days, and liver RNA sequencing was conducted. The heatmap shows DEGs shared by the top KEGG pathways. This suggests that their enrichment may reflect shared mitochondrial gene activation rather than distinct disease-specific processes.

### Supplemental Major Resources Tables

#### Mouse Models (in vivo studies)

| Mouse Model | Vendor or Source | Stock No. | Notes |
|---|---|---|---|
| **C57BL/6J mice** | The Jackson Laboratory | 000664 | Around 12 weeks old; male & female used for all experiments |

#### Mouse Housing Conditions

| Parameter | Mouse Housing Conditions | Note |
|---|---|---|
| **Set Temperature range** | 22 °C | |
| **Set Humidity range** | 50% | |
| **Light Cycle (Mouse)** | 14 hours : 10 hours | |
| **Water** | RO Water | ad libitum |

| | | |
|---|---|---|
| **Standard Feed** | Teklad Irradiated Global 18% Protein Rodent Diet (Envigo); Diet # 2918 | ad libitum |
| **Standard Bedding** | P.J. Murphy Coarse SaniChip | |
| **High Cholesterol Diet Feed** | TD.88051; 7.5% cocoa butter, 15.8% fat, 1.25% cholesterol, 0.5% sodium cholate | ad libitum |
| **SPF** | Yes | |

**Antibodies**

| **Antibody** | **Vendor or Source** | **Catalog #** |
|---|---|---|
| **Anti-CD16/32 (Fc Block)** | BioLegend | 101302 |
| **APC-conjugated anti-CD45** | BioLegend | 103124 |
| **Percp5.5-conjugated anti-CD11b** | BD Biosciences | 550993 |
| **FITC-conjugated anti-Ly-6C** | BioLegend | 128006 |
| **PE-conjugated anti-Ly-6G** | BioLegend | 127608 |
| **B220-FITC** | BioLegend | 103206 |

| | | |
|---|---|---|
| **CD3-PE/Cy7** | BioLegend | 100220 |

**Reagents/Kits/Commercial Service**

| **Reagents** | **Vendor or Source** | **Catalog #** |
|---|---|---|
| **Free Cholesterol E** | Wako | 435-35801 |
| **Total Cholesterol E** | Wako | 439-17501 |
| **Cytokine panel assay** | EVE Technologies | |
| **Corticosterone ELISA Kit** | ENZO Life Sciences | ADI-900-097 |
| **Glucose Meter Kit** | Contour | |
| **Glucose Test Strips** | Contour | |
| **ALT assay kit** | Thermofisher | NC9914003 |
| **BUN assay kit** | BioAssay Systems | DIUR-100 |
| **Rotenone** | Sigma-Aldrich | |
| **30% Acrylamide/Bis solution** | Bio-rad | 161-0156 |
| **APS** | Bio-rad | 161-0700 |

| | | |
|---|---|---|
| **Resolving gel buffer 1.5M Tris-HCL PH 8.8** | Bio-rad | 161-0798 |
| **Stacking gel buffer 0.5M Tris-HCL PH 6.8** | Bio-rad | 161-0799 |
| **TEMED** | Bio-rad | 161-0800 |
| **#23G needle** | Fisher | 14-826A |
| **DPBS** | Sigma | D8662-500 |
| **Fatty acid free BSA** | Sigma | A7030-100g |
| **RNeasy Mini Kit** | QIAGEN | 74104 |
| **RNA-seq service** | Novogene | |
